# A conserved mitochondrial role for cyclin C in mediating oxidative stress-induced cell death in yeast

**DOI:** 10.64898/2025.12.30.696989

**Authors:** Steven J. Doyle, Justin R. Bauer, David C. Stieg, Samar Emami, Daniel G. J. Smethurst, Randy Strich

**Author notes:** Corresponding author, Two Medical Center Drive, 350 Science Center; Rowan School of Osteopathic Medicine, Stratford, NJ 08084. S.J.D. 8565666046, J.R.B. 8565666046, S.E. 8565666046, D.C.S. 8565666046, D.G.J.S.

## Abstract

Cyclin C is a highly conserved component of the Mediator kinase module that regulates transcription through CDK8 stimulation. In addition, the yeast (Cnc1) and human (CCNC) cyclin C exhibit stress-induced mitochondrial translocation to stimulate fission through direct interaction with the dynamin-like GTPase DRP1 (mammals) or Dnm1 (yeast). Gene ablation studies revealed that both Cnc1 and CCNC are required for cell damage-induced regulated cell death (RCD). To determine the relative contributions of cyclin C’s transcriptional and mitochondrial roles in promoting RCD, this study utilized docking simulation algorithms to predict interaction interfaces for CCNC-CDK8 and CCNC-DRP1 heterodimers. As expected, CCNC bound CDK8 through its amino terminal cyclin box domain while DRP1 associated with the second carboxyl cyclin box. Using these predictions, we used site directed mutagenesis on Cnc1 to separate these functions. Importantly, only the DRP1/Dnm1-interaction residues were important for RCD in yeast. Interestingly, although Dnm1 is required for RCD, its fission activity was not. Moreover, Dnm1 is still required for RCD even when Cnc1 is targeted to the mitochondria indicating it is not simply functioning as a tether. Finally, when expressed in yeast, the human CCNC efficiently induced fission and stimulated RCD. Moreover, these functions required predicted DRP1 interaction sites as well. In conclusion, these studies revealed that cyclin C separates its nuclear and mitochondrial activities by utilizing different cyclin box domains. Second, Dnm1-cyclin C interaction, and not transcriptional control, is critical for cyclin C-dependent RCD and this role is conserved from yeast to humans.

**Author Summary:** The cyclin C protein is found in all eucaryotes and exhibits two conserved functions. First cyclin C activates the CDK8 protein kinase to control transcription of genes involved in the stress response. Second, stress induces cyclin C translocation to the mitochondria where it induces fragmentation by stimulating the fission GTPase DRP1. Mitochondrial fission is an early step in the regulated cell death pathway. Importantly, cyclin C is required for stress-induced cell death raising the question of what the relative contribution of its transcription and mitochondrial roles is. To address this question, we generated mutations in the yeast cyclin C that interfered with its association with CDK8 or DRP1 without affecting the other. We found that the mitochondrial role, but not transcription, was required for cell death in yeast. In addition, when expressed in yeast, the human cyclin C also induced mitochondrial fission and cell death. Finally, although DRP1 is important for regulated cell death, its fission function is not. These results point to a highly conserved role for cyclin C in mediating regulated cell death at the mitochondria. In addition, these results point to a non-enzymatic role for DRP1 in executing the cell death pathway.

## Introduction

Surviving cellular damage requires a coordinated response between the various organelles. For example, changes in the nuclear transcriptome profile increases stress response gene transcription while reducing the expression of those loci involved in cell division (1). Simultaneously, the mitochondria undergo extensive fission that prepares the cell for survival (e.g., mitophagy) or intrinsic regulated cell death (iRCD) (2). Which cell fate is taken depends on many parameters including the duration and extent of damage incurred.

The cyclin protein family binds and activates cyclin-dependent protein kinases (Cdks) responsible for driving cell cycle progression through substrate phosphorylation (3). Cyclins were named based on their periodic expression pattern during the cell cycle (4). A second class of cyclins that primarily control transcription exhibits fixed expression through the cell cycle (5, 6). Cyclin C and its cognate kinases Cdk8 or Cdk19, are components of the Mediator Kinase Module (MKM) that play a direct role controlling transcription through its association with the Mediator complex (7). The MKM acts primarily as a transcriptional repressor in yeast (8, 9) but exhibits both repressor and activator functions in mammalian cells (10).

We previously reported that both the yeast (Cnc1) and mammalian (CCNC) cyclin C family members play dual roles in mediating the stress response. In the nucleus, both Cnc1 and CCNC repress genes upregulated during oxidative stress and are required for full induction of other stress responsive genes (10, 11). In addition, oxidative stress also triggers re-localization of Cnc1 or CCNC to the cytoplasm (12–14). In the cytoplasm, both Cnc1 and CCNC associate with the mitochondrial fission proteins Mdv1-Dnm1 (yeast) (12) or DRP1 (mammals) (15) which in turn stimulates mitochondrial fission and helps prepare the cell for RCD in response to stressors such as prooxidants (e.g., H_2_O_2_) or anti-cancer therapies (e.g., cisplatin) (16). In mammalian cells, CCNC binds the proapoptotic protein Bax recruiting it to the mitochondrial outer membrane (MOM) to promote mitochondrial outer membrane permeability (MOMP), an early decision point for RCD in metazoans (17). MOMP promotes the release of cytochrome *c* which in turn stimulates caspase activation and ultimately cell death. However, the absence of Bcl-2 proteins in yeast raises the question of how the mitochondria support RCD.

Given the conserved role of cyclin C in both the nucleus and mitochondria, the relative contributions of each activity in promoting RCD were unknown. Cyclins were initially defined by the signature motif termed the cyclin box (CB), a domain composed of a five alpha helical bundle that binds the Cdk (18). Interestingly, many cyclins, including cyclin C, have amino and carboxyl cyclin boxes here termed CB1 and CB2, respectively. While CB1 directs Cdk binding in all cyclins tested, we demonstrated that CB2 is necessary and sufficient for binding Drp1 in vitro and stimulating its activity in cellular assays. These results suggest a modular organization for cyclin C in that the individual functions are executed by different CB domains. This model raises the possibility that individual Ccnc functions can be separated by mutating the domain directing one function or the other. This report used prior crystal structure studies, alpha fold predictions and docking algorithms to predict CDK8 and DRP1 interacting amino acids on CCNC. Using this information, mutations were generated in the yeast Cnc1 that disrupted its activity with Dnm1 or Cdk8 and their effect on RCD determined. These studies revealed the Dnm1-Cnc1 interaction was critical for RCD execution but not Cnc1-Cdk8. However, although Dnm1 is required for cyclin C-dependent cell death, its fission function is not. Finally, the human CCNC provides mitochondrial and RCD activity in yeast demonstrating the CB2 function is highly conserved within this cyclin family.

## RESULTS

### Validating the structural analysis tools

Our previous studies revealed that cyclin C possesses conserved roles in stimulating mitochondrial fission and regulating transcription by directly binding DRP1 and CDK8, respectively. To identify potential interaction regions between cyclin C and DRP1 or CDK8, we sought conserved domains between the yeast (Cnc1) and human (CCNC) proteins. First, we examined alignments of both modeled and experimentally derived structures of the two proteins. Using PyMol’s built in alignment tool (19), a high degree of structural homology was observed between experimentally solved yeast Cnc1 (PDB 7KPX652) and human CCNC (PDB 5CEI654) structures (Fig S1A). Importantly, the five helix bundle regions containing the CB1 and CB2 domains are highly conserved. To quantify the structural similarities, root mean square deviation (RMSD) scores (among all atoms) determined a deviation between 1.289-4.221 (Fig S1B), depending on the distance cutoff. These results indicate that the overall three-dimensional structures of the yeast and human proteins are conserved.

We next compared PyMol predictions of the cyclin C-Cdk8 heterodimer using experimentally derived structures and the ClusPro protein-protein docking simulation algorithm (20–23). ClusPro operates in a three-step computation process to identify favorable charge and hydrophobic interactions between potential docking interfaces (see Methods for details). To account for the residues not included in the experimental structures, AlphaFold protein-modeling was used to provide the remaining domains (24, 25). When compared to the experimentally derived structure coordinates, the best scoring ClusPro model accurately predicted the heterodimer structure (Fig S1C). As expected, poorer scoring models produced higher RMSD values when compared to the experimentally derived structure (Fig S1D). Taken together, these results indicate that the ClusPro algorithm produces accurate heterodimer models that have not been solved experimentally.

### Identifying the CCNC-CDK8 interaction interface

To determine the relative contributions of the two cyclin C functions on iRCD, we sought to generate mutant alleles that abolished either its transcription or mitochondrial function. To accomplish this, we sought residues defining the interactions between CCNC-CDK8 or CCNC-DRP1. We utilized the human proteins as more examples of the solved crystal structures were available. Using ClusPro modeling, the docking interface for the CCNC-CDK8 heterodimer was analyzed first. Using a cutoff frequency of 20% inclusion for interacting amino acids in the top 100 scoring models, we identified 19 models for further evaluation. The top scoring frequency of a specific amino acid interaction in all models generated was 33%, referred to as the “hit%” (Table 1). The predicted type of interaction, hydrogen bonding and/or salt bridge formation, are indicated. The CCNC domain containing the interaction residues was consistently limited to CB1 particularly alpha helix 3 and 5 (Fig 1A, see open/blue flags, Fig 1C). These results are consistent with CCNC-CDK8 interactions described in crystal (18) or cyro-EM (26) structural analyses (Fig S1C and Table 1). ClusPro modeling identified both CCNC-CDK8 residue pairs either identical to experimentally derived structures or only identified by the algorithm. For the top 14 predictions listed, nine matched both CCNC and CDK8 residues (Fig 1A, mapped in Fig 1C). For example, the cryo-EM structure and the ClusPro docking simulations identified similar residues (K96, E99, E137, E144) that direct CCNC-CDK8 interaction (Fig 1A, open/blue flags, Fig 1C). In addition, several of the CCNC amino acids were also predicted to interact with additional CDK8 amino acids. For example, Glu144 was predicted to form a salt bridge with Arg12 of CDK8, an interaction observed in the crystal structure. However, Glu144 was also predicted to form hydrogen bond with Lys185. This pattern was observed for other CCNC amino acids as well. Taken together, ClusPro modeling identified bone fide interactions but also suggested additional interactions at similar frequencies.

**Figure 1.**
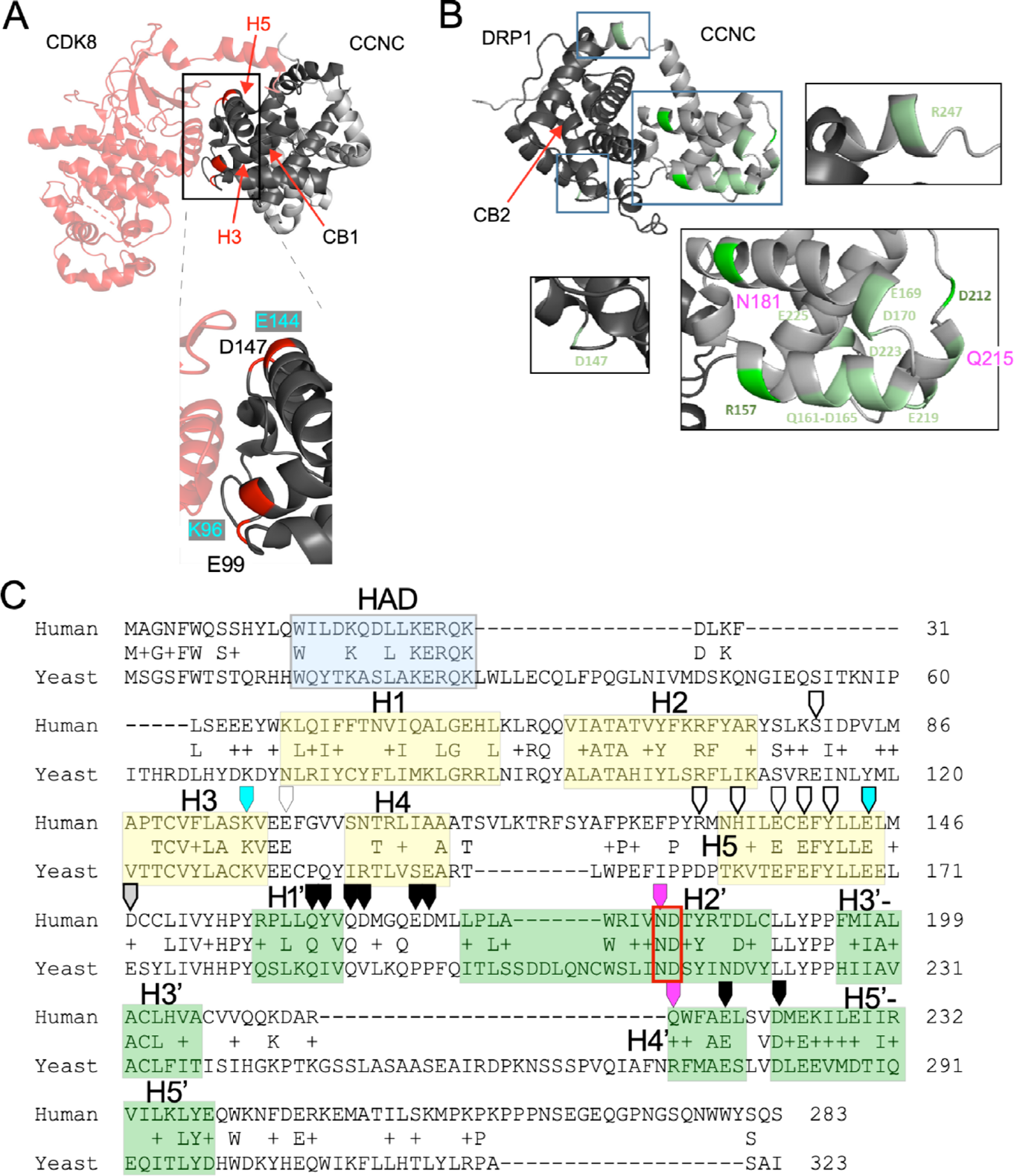
Predicted CDK8 and DRP1 interaction regions of CCNC. ClusPro modeling of CCNC-CDK8 (A) and CCNC-DRP1 (B) predicted interactions. Boxes indicate exploded regions for both panels. Residues of interest are indicated. Blue (K96, E99) and fuchsia (N181, Q215) are studied further in detail. (C) Human and yeast cyclin C sequences are shown. Identical amino acids and conservative substitutions (+) are indicated between the sequences. Yellow boxes denote the helices (H1-H5) associated with cyclin box 1 (CB1) and green boxes the five helices (H1’-H5’) defining CB2. Open flags indicate CCNC-CDK8 interactions with the blue flags highlighting K96 and E144. Closed flags indicate predicted CCNC-DRP1 interactions. Fuchsia flags denote N181 and Q215. Grey flag (Asp 147) indicates only amino acid predicted to interact with both CDK8 and DRP1. Blue box indicates the holoenzyme association domain (HAD) required for MED13 interaction (see text for details). Red box indicates amino acids targeted for CCNC mutagenesis.

**Table 1.** Comparison of interacting residues of ClusPro CCNC-CDK8 binding simulations and experimentally derived structure. Hit % indicates percentage of all models with residue pair (CCNC/CDK8) at the interaction interface. ^a^type identifiers include hydrogen bond and/or salt bridge formation. Predictions that match the experimentally derived structure at both the CCNC and CDK8 residues are indicated. CCNC only indicates a model that identifies only a CCNC interacting residue but not the experimentally determined CDK8 amino acid. Bold face type indicates two amino acids targeted for further study.

| CCNC |  | CDK8 | Interaction Type | Models |  |
| --- | --- | --- | --- | --- | --- |
| AA | Domain | AA | Type<sup>a</sup> | Hit % | Matches Crystal Structures |
| Glu144 | $\alpha 5$ | Arg13 | Hydrogen/Salt | 33% | YES |
| Glu144 | $\alpha 5$ | Lys185 | Hydrogen | 33% | CCNC only |
| Lys96 | $\alpha 3$ | Cys64 | Hydrogen | 27% | CCNC only |
| Asp147 | $\alpha 5$ | Arg71 | Hydrogen/Salt | 27% | YES |
| Ser80 | $\alpha 2/\alpha 3$ | Asp2 | Hydrogen | 20% | YES |
| Lys96 | $\alpha 3$ | Ile59 | Hydrogen | 20% | YES |
| Glu99 | $\alpha 3/\alpha 4$ | Arg91 | Hydrogen/Salt | 20% | CCNC only |
| Glu99 | $\alpha 3/\alpha 4$ | Arg178 | Salt | 20% | YES |
| Glu99 | $\alpha 3/\alpha 4$ | Lys185 | Hydrogen/Salt | 20% | CCNC only |
| Leu136 | $\alpha 5$ | Arg91 | Hydrogen | 20% | YES |
| Glu137 | $\alpha 5$ | Lys6 | Hydrogen/Salt | 20% | YES |
| Glu137 | $\alpha 5$ | His88 | Salt | 20% | CCNC only |
| Lys261 | C-term | Asp2 | Hydrogen | 20% | YES |
| Lys261 | C-term | Asp4 | Hydrogen/Salt | 20% | YES |

as DRP1 interactors (see Table 2). A cluster of predicted interacting amino acids was also observed in the region containing H4’ and H5’. These interactions consisted largely of charged amino acids. One exception was a strong predicted interaction site at N181 in H2’ (Fig 1B and pink flag, Fig 1C). Taken together, these results predict the H1’-H2’ and H4’-H5’ regions as CCNC-DRP1 interaction sites. However, additional contacts in H2’ were also indicated.

**Table 2.** Predicted CCNC-DRP1 interaction sites. CCNC residues and their predicted DRP1 contacts are listed. Interaction types are included as hydrogen binding or salt bridge formation. %Hit indicates the percentage of top 100 models included the individual CCNC residue with potential DRP1 residues indicated.

| CCNC |  |  | DRP1 |  |  | Interaction Types | Models |
| --- | --- | --- | --- | --- | --- | --- | --- |
| AA | AA# | Domain | Domain | AA# | AA | Type<sup>a</sup> | Hit % |
| Asp | 147 | $\alpha 5/\alpha 1'$ | GTPase | 141 | Asn | Hydrogen | 45% |
|  |  |  |  | 208 | Arg | Hydrogen/Salt |  |
|  |  |  |  | 238 | Lys | Hydrogen/Salt |  |
|  |  |  |  | 278 | Asn | Hydrogen/Salt |  |
|  |  |  |  | 279 | Arg | Hydrogen/Salt |  |
|  |  |  |  | 283 | Lys | Hydrogen/Salt |  |
|  |  |  |  | 287 | Arg | Hydrogen/Salt |  |
|  |  |  |  | 290 | Asn | Hydrogen |  |
| Asp | 165 | $\alpha 1'/\alpha 2'$ | GTPase | 278 | Asn | Hydrogen | 40% |
|  |  |  |  | 279 | Arg | Salt |  |
|  |  |  |  | 287 | Arg | Hydrogen/Salt |  |
|  |  |  |  | 291 | Arg | Hydrogen/Salt |  |
|  |  |  |  | 305 | Lys | Hydrogen/Salt |  |
| Glu | 219 | $\alpha 4'$ | GTPase | 279 | Arg | Hydrogen/Salt | 35% |
|  |  |  |  | 280 | Asn | Hydrogen |  |
|  |  |  |  | 284 | Tyr | Hydrogen |  |
|  |  |  |  | 291 | Arg | Hydrogen/Salt |  |
|  |  |  |  | 295 | His | Salt |  |
|  |  |  |  | 298 | Arg | Hydrogen/Salt |  |
|  |  |  |  | 305 | Lys | Hydrogen/Salt |  |
| Gln | 161 | $\alpha 1'/\alpha 2'$ | GTPase | 69 | His | Hydrogen | 25% |
|  |  |  |  | 73 | Glu | Hydrogen |  |
|  |  |  |  | 141 | Asn | Hydrogen |  |
|  |  |  |  | 291 | Arg | Hydrogen |  |
|  |  |  |  | 298 | Arg | Hydrogen |  |
|  |  |  |  | 303 | Glu | Hydrogen |  |
| Glu | 169 | $\alpha 1'/\alpha 2'$ | GTPase | 69 | His | Salt | 25% |
|  |  |  |  | 72 | Gln | Hydrogen |  |
|  |  |  |  | 305 | Lys | Salt |  |
| Gln | 215 | $\alpha 4'$ | GTPase | 230 | Leu | Hydrogen | 15% |
|  |  |  |  | 279 | Arg | Hydrogen |  |
|  |  |  |  | 305 | Lys | Hydrogen |  |
| Asp | 223 | $\alpha 5'$ | GTPase | 208 | Arg | Salt | 15% |
|  |  |  |  | 291 | Arg | Hydrogen/Salt |  |
|  |  |  |  | 295 | His | Salt |  |
|  |  |  |  | 298 | Arg | Hydrogen/Salt |  |
| Gln | 164 | $\alpha 1'/\alpha 2'$ | GTPase | 137 | Pro | Hydrogen | 15% |
|  |  |  |  | 141 | Asn | Hydrogen |  |
|  |  |  |  | 290 | Asn | Hydrogen |  |
| Asp | 170 | $\alpha 2'$ | GTPase | 287 | Arg | Hydrogen/Salt | 15% |
|  |  |  |  | 305 | Lys | Hydrogen/Salt |  |
| Tyr | 162 | $\alpha 1'/\alpha 2'$ | GTPase | 291 | Arg | Hydrogen | 15% |
|  |  |  |  | 298 | Arg | Hydrogen |  |

### Identifying the CCNC-CDK8 interaction interface

To determine the relative contributions of the two cyclin C functions on iRCD, we sought to generate mutant alleles that abolished either its transcription or mitochondrial function. To accomplish this, we sought residues defining the interactions between CCNC-CDK8 and CCNC-DRP1. We utilized the human proteins as more examples of the solved crystal structures were available. Using ClusPro modeling, the docking interface for the CCNC-CDK8 heterodimer was analyzed first. Using a cutoff frequency of 20% inclusion for interacting amino acids in the top 100 scoring models, we identified 19 models for further evaluation. The top scoring frequency of a specific amino acid interaction in all models generated was 33%, referred to as the “hit%” (Table 1). The predicted type of interaction, hydrogen bonding and/or salt bridge formation, are indicated. The CCNC domain containing the interaction residues was consistently limited to CB1 particularly alpha helix 3 and 5 (Fig 1A, see open/blue flags, Fig 1C). These results are consistent with CCNC-CDK8 interactions described in crystal (18) or cyro-EM (26) structural analyses (Fig S1C and Table 1). ClusPro modeling identified both CCNC-CDK8 residue pairs either identical to experimentally derived structures or only identified by the algorithm. Several CCNC-CDK8 interaction pairs predicted by the algorithm were also identified in the crystal structure. For the top 14 predictions listed, nine matched both CCNC and CDK8 residues (Fig 1A, mapped in Fig 1C). For example, the cryo-EM structure and the ClusPro docking simulations identified similar residues (K96, E99, E137, E144) that direct CCNC-CDK8 interaction (Fig 1A, open/blue flags, Fig 1C). In addition, several of the CCNC amino acids were also predicted to interact with additional CDK8 amino acids. For example, Glu144 was predicted to form a salt bridge with Arg12 of CDK8, an interaction observed in the crystal structure. However, Glu144 was also predicted to form hydrogen bond with Lys185. This pattern was observed for other CCNC amino acids as well. Taken together, ClusPro modeling identified bone fide interactions but also suggested additional interactions at similar frequencies.

### Identifying the CCNC-DRP1 interaction interface

All the experimentally determined DRP1 structures solved the GTPase domain and omitted a large region of the the C-terminus. Using PrDos (27) and flDPnn (28) analysis tools, we confirmed the presence of a large intrinsic disordered region (IDR) in this region and omitted it from our analysis (see Fig S2). In addition, our prior results indicated that the GTPase domain (residues 1-340) was necessary and sufficient for CCNC interaction (15). We chose the starting structural inputs of CCNC (PDB 6Y0A) and DRP1 (PDB 4H1V). Consistent with our previous results, ClusPro docking predictions indicated that nearly all DRP1 GTPase domain interactions occurred within CB2 (Fig 1B and 1C). Specifically, the charged (D165, E169, D170) and hydrophobic/polar (V163, Q164, Q168) amino acids between helices H1’ and H2’ scored high as DRP1 interactors (see Table 2). A cluster of predicted interacting amino acids was also observed in the region containing H4’ and H5’. These interactions consisted largely of charged amino acids. One exception was a strong predicted interaction site at N181 in H2’ (Fig 1B and pink flag, Fig 1C). Taken together, these results predict the H1’-H2’ and H4’-H5’ regions as DRP1 interaction sites. In addition, contacts in H2’ were also indicated. Taken together, these results suggest that the two activities of cyclin C have been partitioned into CB1 and CB2.

### DRP1 interaction sites are required for Cnc1-dependent fission in yeast

We chose the yeast system to rapidly test the functionality of potential CDK8 and DRP1 interaction amino acids in directing mitochondrial fission. This approach was based on our finding that recombinant yeast Cnc1, when added to permeabilized MEF cells, can rapidly induce mitochondrial fragmentation (13) indicating functional conservation for this activity. First, the homologous amino acids identified as potential interactors on CCNC were mapped to Cnc1 (Fig S3A). As expected, CDK8 and DRP1 interaction sites also mapped to CB1 and CB2, respectively (see Fig S3B). Alanine substitution mutations were introduced in a yeast *CNC1*-YFP reporter gene. In addition, this reporter was fused to the C-terminal mitochondrial outer membrane transmembrane domain of the Dnm1 receptor Fis1 (*CNC1*-YFP-Fis1) (29, 30). We previously reported that the Cnc1-YFP-Fis1 reporter targets this protein to the mitochondria and induces fission in the absence of stress signals (31). In log-phase unstressed wild-type and *cnc1Δ* cells, mitochondria exhibit an elongated reticular morphology (Fig 2A, left and middle panels). As we previously reported (31), Cnc1-YFP-Fis1 localizes to the mitochondria (yellow foci) inducing fission in ∼88% of the population (right panel). First, alanine substitutions were introduced into residues predicted to bind CDK8. We chose alanine substitutions in hopes of minimizing any negative impact on Cnc1’s mitochondrial function directed by CB2. Alanine substitutions were introduced at K130, E133 or E162 (Fig 2B). All Cnc1-YFP-Fis1 derivatives were expressed and associated with the mitochondria as determined by fluorescence microscopy. Mitochondrial fission occurred at wild-type levels in cells expressing the K130A (96%) and E162A (80%) derivatives indicating that these mutant proteins still stimulate Dnm1 activity. However, the E133A mutation exhibited an intermediate value (27%) (Fig 2B, middle panel). We examined three additional mutations (Y166A, E169A and E172A) that targeted CDK8 interacting residues. All three mutant proteins were able to execute mitochondrial fission at wild-type efficiency (Fig 2C). These results indicate that predicted CDK8 interaction mutants were still able to stimulate mitochondrial fission.

**Figure 2.**
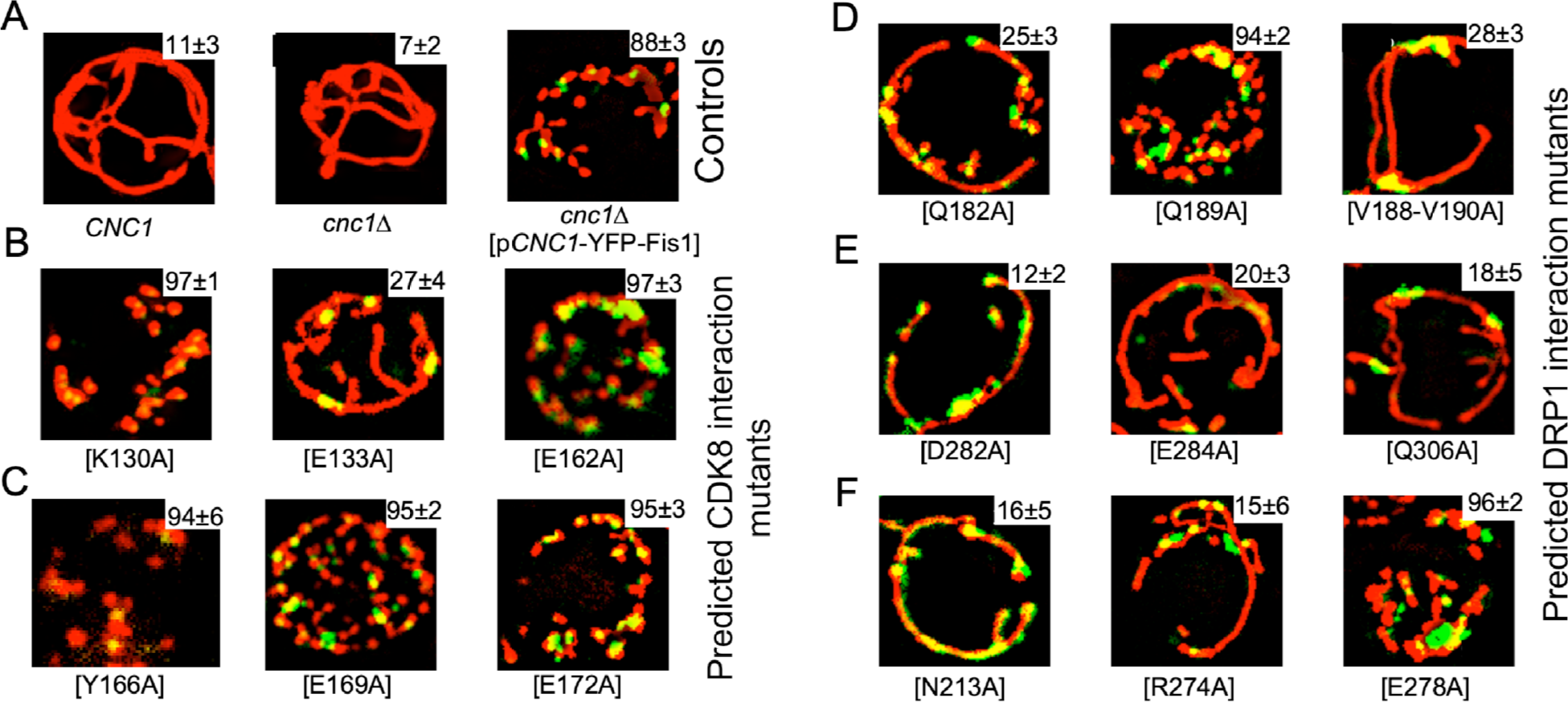
Fission assays for predicted CDK8 and DRP1 interaction mutants. (A) Wild type, *cnc1Δ* and *cnc1Δ* cells transformed with a mitochondrial targeted YFP-*CNC1*-*Fis1* reporter expression plasmids with the mutations indicated. Mitochondria were visualized using MitoTracker Red. Mitochondrial fragmentation ability of the different *CNC1* derivatives are shown in the upper right corner of each panel (see methods for details on quantitation). (B) and (C) The *cnc1Δ* mutant strain was transformed with the same expression plasmid harboring mutations in predicted CDK8 interaction residues as indicated. (D), (E) and (F). Same as (B) except these cells expressed *CNC1* alleles mutated for potential Dnm1 interaction sites as indicated. See Fig S3A for the summary of these results and corresponding yeast Cnc1 amino acids.

Next, we generated alanine substitutions at nine amino acids predicted to interact with Dnm1. Q182A reduced fission (20% fragmented mitochondria) while the Q189A had no effect (87%) (Fig 2D, left and center panels). However, when the two residues flanking Q189 were also mutated (V188A-Q189A-V190A), a significant reduction in fission was observed (Fig 2D, right panel). Taken together, these results indicate that the Cnc1-Dnm1 interactions directed through these hydrophobic/non-polar amino acids are required for normal mitochondrial fission. Finally, residues predicted to interact with Dnm1 in H2’ (N213A) and H4’-H5’ (R274A, D282A and E284A) were also defective for fission (Figs 2E and 2F). Interestingly, mutating Q306, a residue predicted to interact with DRP1/Dnm1 but lies outside the predicted interaction face (Fig S3B), still allowed Cnc1-YFP-Fis1 fission. These results identify an interaction domain for DRP1/Dnm1 that resides in CB2 within the H1’-H2’ and H4’-H5’ regions. In addition, these studies suggest a strong evolutionary conservation as these regions maintain strong identity between yeast and human proteins.

### Cdk8 interaction amino acids are required for transcriptional repression

Next, we tested the transcriptional functionality of the CDK8 and DRP1 interaction mutants again using the yeast system. First, we identified two analogous residues K130 (H3) and E133 (H5) in the yeast Cnc1 corresponding to the CCNC K96 and E99 interaction sites with CDK8 (Fig 1C, listed in Fig 3A). Similarly, we chose N213A (H2’) and R274A (H5’) fission mutants to test in the transcription assays. To test the transcriptional role of these residues, alanine substitutions were generated at each site in a myc-*CNC1*-YFP reporter gene. The Fis1 domain was removed to allow nuclear localization of these derivatives. As Cnc1 predominantly represses transcription in yeast (9), we chose two genes (*DDR2* and *HSP26*) whose transcription is repressed by Cnc1-Cdk8 (32). Ddr2 is involved in multiple stress responses (33) while Hsp26 is a chaperone that prevents misfolded protein aggregation (34). RT-qPCR was performed on total RNA preparations in *cnc1Δ* mutant cells harboring expression plasmids expressing the individual mutant genes. Compared to wild-type control, the *cnc1Δ* strain harboring the vector alone exhibited a five-fold increase in *DDR2* mRNA levels (Fig 3B). The K130A expressing cells also displayed elevated *DDR2* expression similar to the *cnc1Δ* strain. The E169A mutation also significantly derepressed *DDR2* transcription although to a reduced level compared to the K130A allele. The two Dnm1 interacting mutant proteins, N213A and R274A, continued to repress *DDR2* transcription to wild-type levels. A similar result was obtained for *HSP26* mRNA levels (Fig 3C). These results indicate that the predicted Cdk8 interaction animo acids are important to maintain transcriptional repression while the Dnm1 interaction residues are dispensable.

**Figure 3.**
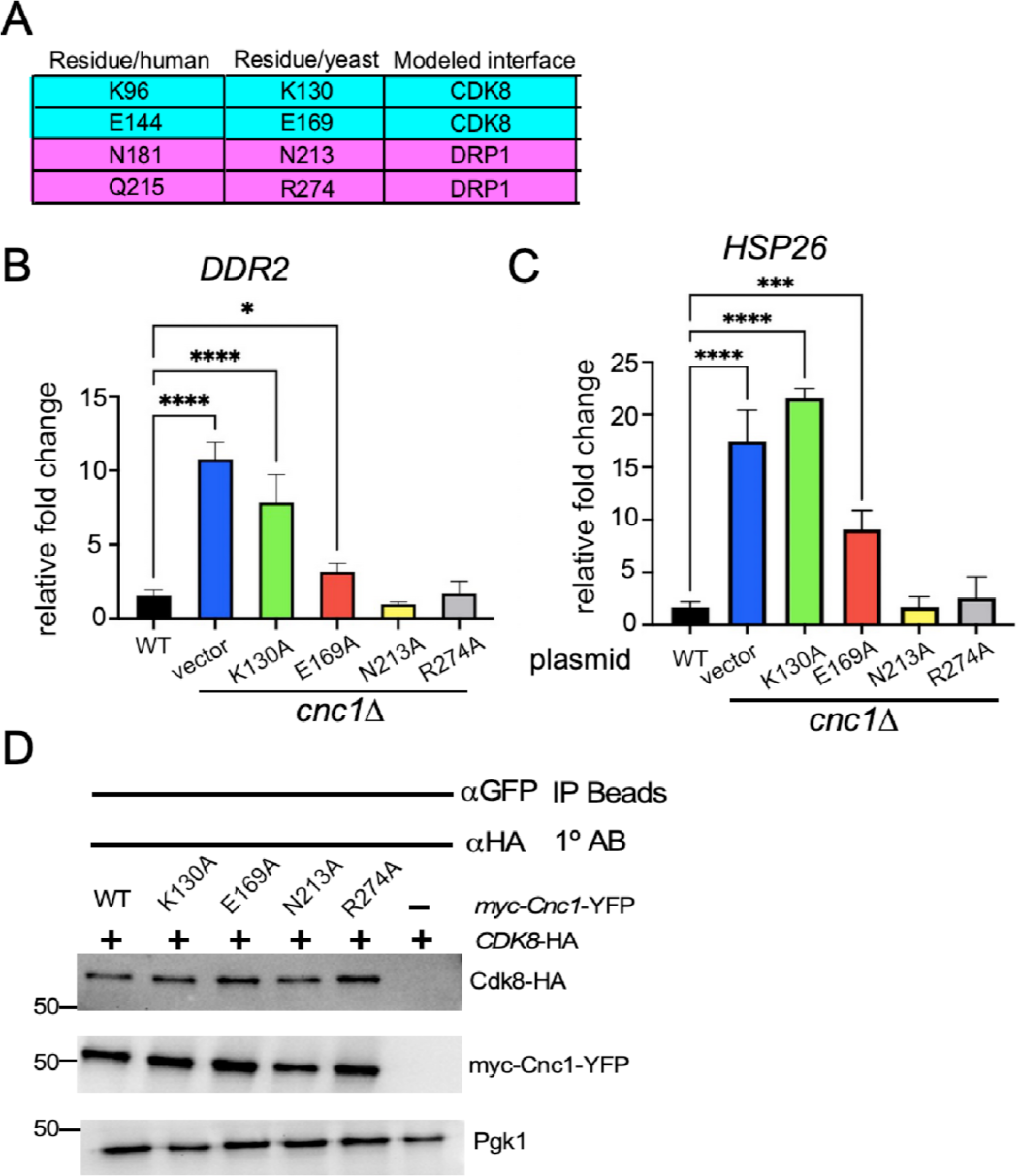
Transcriptional repression assays for predicted CDK8 and Dnm1/DRP1 interaction mutants. (A) Conversion table for human to yeast mutant amino acids used in this study including the modeled protein partner. (B) and (C) RT-qPCR analysis of *DDR2* and *HSP26* mRNA levels in wild type cells and the *cnc1Δ* mutant harboring either the vector alone or expression constructs (myc-*CNC1-*YFP) carrying the indicated mutations. *ACT1* mRNA was used as the mRNA control for the ΔΔc_t_ calculations (N = 3 biological replicates with three technical replicates each). Asterisks *, p=0.05, ***, p=0.01, **** p=0.001. (D) Co-immunoprecipitation assays were performed in extracts prepared from cells harboring the indicated expression plasmids. The extracts were immunoprecipitated with αGFP beads prior to Western blot analysis probing for the HA epitope. This blot was stripped then probed for the myc epitope to visualize Cnc1-YFP. Pgk1 levels were monitored by Western blot analysis of a separate gel as a control for extract quantification. Molecular weight markers (kDa) are indicated.

One possible mechanism to explain these results is that the K130A and the E133A mutations disrupted interaction with Cdk8. To address this question, co-immunoprecipitation experiments were performed (see Materials and methods). Soluble protein extracts were prepared from log phase cultures each harboring plasmids expressing the individual myc-*CNC1-*YFP derivatives and another expressing the HA epitope tagged *CDK8* (Cdk8-HA). As a control, an extract was also prepared from a *cnc1Δ* strain lacking the myc-*CNC1*-YFP wild-type expression plasmid. These extracts were incubated with αGFP beads to bind the YFP tag. The beads were washed, boiled in sample buffer and subjected to Western blot analysis probing with HA antibodies to detect Cdk8-HA. The results indicated that the mutant proteins were expressed at levels comparable to the wild type (Fig 3D). Importantly, Cdk8-HA was immunoprecipitated at similar efficiencies for the wild type and all Cnc1-YFP derivatives. These results indicate that the association of Cnc1 with Cdk8 is not reduced by the K130A or E169A mutations. However, given the overall structure of the MKM, it is possible that Cnc1-Cdk8 activity is altered but that other contacts within this complex still allow co-immunoprecipitation (see Discussion).

### Dnm1 interaction is required for Cnc1-dependent stress-induced mitochondrial fission

Our earlier experiments indicated that mitochondrial targeted Cnc1 derivatives mutated at Dnm1 interaction sites failed to induce fission. However, this artificial construct did not undergo nuclear to mitochondrial translocation following cellular stress. To better understand the impact these mutations had on mitochondrial fission, we monitored myc-Cnc1-YFP localization and mitochondrial fragmentation following oxidative stress. In wild-type cells, myc-Cnc1-YFP was nuclear in unstressed cells but rapidly localized to the mitochondria following a 2 h exposure to 50 µM cumene hydroperoxide (CHP, Fig 4A) resulting in >80% of the cells exhibiting extensive fission. The same Cdk8 interaction mutants just described (K130A and E169A) were expressed in a *cnc1Δ* mutant strain. Following CHP exposure, both transformants displayed mitochondrial re-localization and wild-type levels of fragmentation (Figs 4B and 4C). Interestingly, the Dnm1 interaction mutants (N213A, R274A) displayed normal nuclear localization before CHP treatment and mitochondrial translocation after treatment (Fig 4D and 4E). Despite localizing to the mitochondria, these mutants failed to stimulate fission. These results indicate that the nuclear and mitochondrial roles of Cnc1 can be separated in yeast.

**Figure 4.**
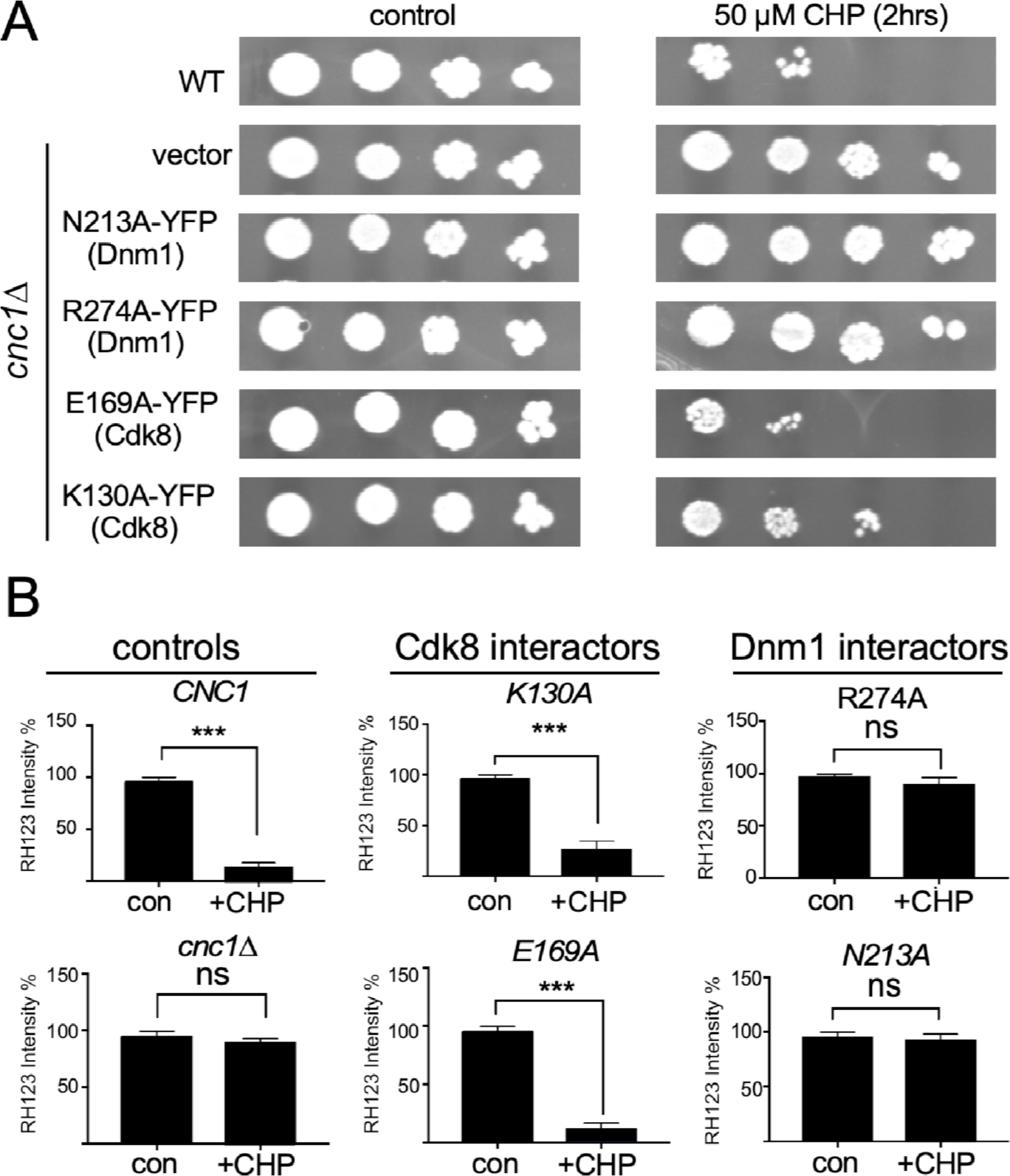
Analysis of nuclear to mitochondrial translocation. (A) Mitochondrial morphology and myc-CNC1-YFP localization were monitored in a *cnc1Δ* null strain before and after CHP treatment (50 µM, 2 h). Percentage of the cells exhibiting fragmented mitochondria was quantified (N=3). (B-E) The experiments in (A) were repeated in *cnc1Δ* cells expressing the indicated *CNC1* mutants. ****, p=0.001, ns = not significant. (F) Extracts from cultures expressing the indicate myc-CNC1-YFP derivatives and Dnm1-myc were immunoprecipitated with YFP then probed from myc. The signals were combined with those obtained from a second trial then the Cnc1/Dnm1 ratios calculated (see Fig S4). The transcription or fission mutant lanes were combined to add statistical significance. The WT value was set at 1.0.

We next examined the ability of the myc-Cnc1-YFP mutants to interact with Dnm1 in stressed cells. The experiments just described were repeated except that a functional *DNM1*-myc expression plasmid was introduced to the strains. The cells were grown to mid-log phase, treated with CHP for four hours, then extracts prepared. Since both Dnm1 and Cnc1 were tagged with myc, both proteins were visualized with this approach. Quantifying the signals from each protein, a Cnc1/Dnm1 ratio was calculated with the wild-type myc-Cnc1-YFP/Dnm1 set to 1.0 (Fig 4F). The Cdk8 mutants, K130A and E169A, immunoprecipitated with Dnm1-myc at wild-type efficiencies. However, extracts derived from cells expressing either N213A or R274A mutants displayed a reduced co-immunoprecipitation ability. Combining these values with another trial (Fig S4A, quantitated in S4B) revealed a significant reduction in myc-Cnc1^K213A^-YFP or myc-Cnc1^R274A^-YFP association with Dnm1-myc. These results indicate that the N213A and R274A mutations reduced, but do not eliminate, Cnc1-Dnm1 interaction.

### Cnc1-Dnm1 interaction, but not mitochondrial fission, is required for CHP-induced cell death

The previous results indicate that we successfully separated the transcription and mitochondrial fission activities of Cnc1. To determine if one, or both functions, are required for normal RCD execution, *cnc1Δ* mutant yeast cells expressing the four mutants described above were treated with CHP for 2 h and viability determined. Our positive and negative controls were the wild-type and *cnc1Δ* cells harboring the vector alone. As expected, the wild-type cells were sensitive to CHP treatment while the *cnc1Δ* cells were not (Fig 5A). Cells expressing the two Cdk8 mutants, E169A and K130A, were also sensitive to CHP treatment. However, both DRP1/Dnm1 mutants, N213A and R274A, were resistant to oxidative stress similar to the *cnc1Δ* cells. These studies indicate that normal Cnc1-Dnm1 association is required for ROS-induced RCD in yeast.

**Figure 5.**
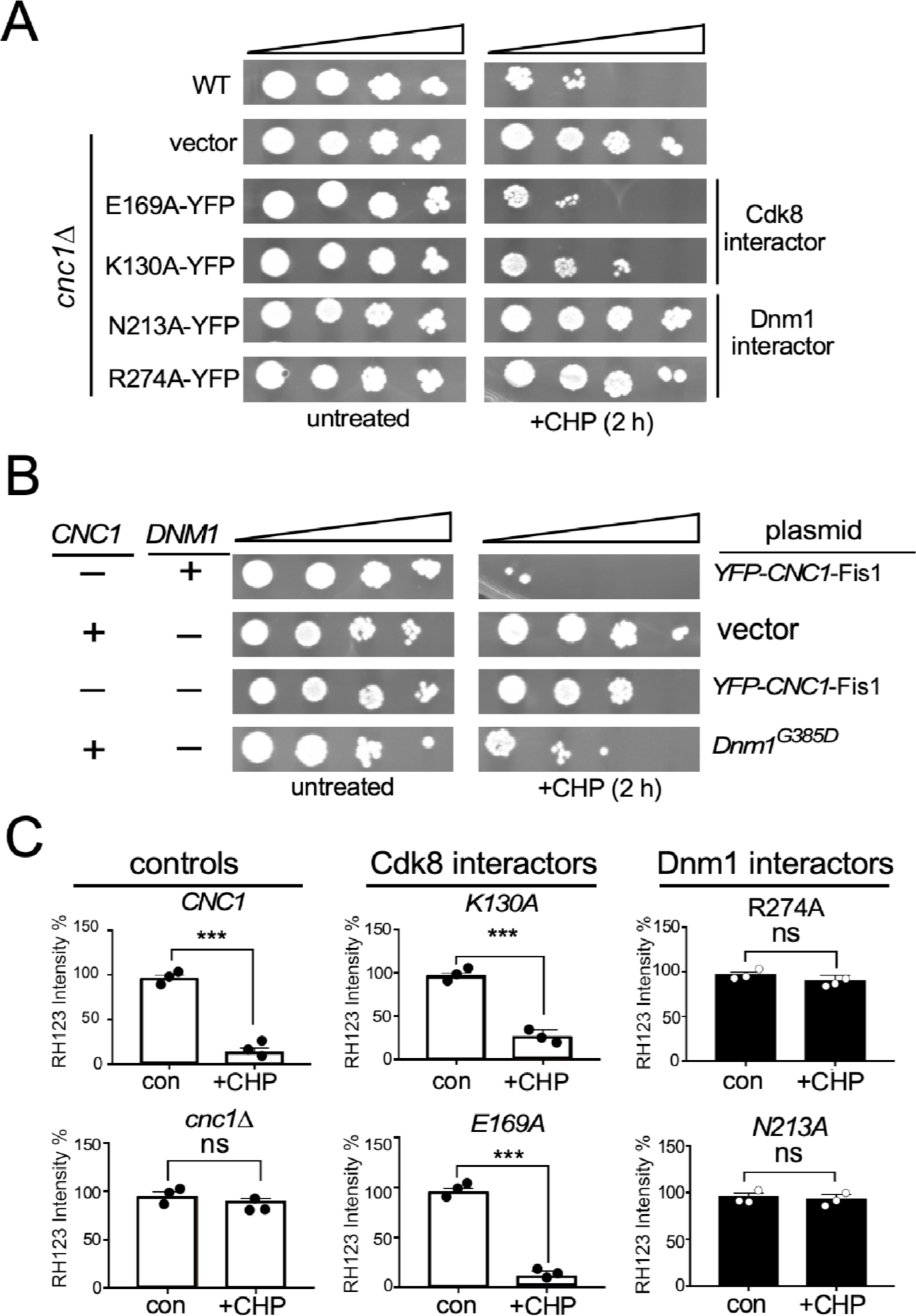
Dnm1 interaction, but not fission, is required for RCD. (A) WT or *cnc1Δ* mutant strains expressing the indicated *CNC1* alleles were grown to mid log, the cultures split, and half treated with CHP. The cultures were serially diluted (1:10) then plated on solid medium. The plates were incubated 2 d then images acquired. (B) Cultures with the indicated genotype harboring expression plasmids listed were treated as described in (A). (C) The experiments described in (A) were repeated except cells were treated with the fluorophore RH123. These cultures were then analyzed by fluorescent cell analysis. *** p=0.005, ns = not significant.

To more fully examine the role of Dnm1 in Cnc1-induced RCD, we used the mitochondrial targeted YFP-Cnc1-FIS1 reporter gene. This construct renders cells hypersensitive to CHP treatment (Fig 5B, top row). As previously reported (35, 36), Dnm1 is required for CHP-induced RCD (Fig 5B, second row). Moreover, Dnm1 is required for cell death even in the presence of the mitochondrial targeted Cnc1-YFP-Fis1 (third row). These results indicate that Dnm1 is performing a function other than simply recruiting Cnc1 to the mitochondria. To test the role of fission itself in this cell death process, we repeated these experiments with a *DNM1* mutation (*Dnm1^G385D^*) that can form dimers but not oligomers capable of fission (37). Cells expressing the *Dnm1^G385D^*allele were still sensitive to CHP treatment indicating the Dnm1, but not Dnm1-induced fission, is required for Cnc1 stimulated cell death. Therefore, Dnm1 is an active participant in promoting cell death although fission itself is not required.

In mammalian iRCD studies, mitochondrial outer membrane permeability (MOMP) represents a critical point in the cell death process. We previously observed that CCNC is required for MOMP in cisplatin treated mammalian cells (13). To determine whether MOMP occurs during the studies described above, we employed the mitochondrial fluorophore RH123. RH123 is excited in the oxidative environment of the mitochondria but loses its fluorescence when released into the cytosol. This change in fluorescence was monitored by fluorescence activated cell analysis. As controls, MOMP was monitored in wild type and *cnc1Δ* cells following CHP treatment. These studies revealed a significant loss in RH123 fluorescence in wild-type cells treated with CHP while fluorescence remained elevated the *cnc1Δ* strain (left panels Fig 5C, see Fig S5 for representative traces). To determine if MOMP occurred in the presence of the Cdk8 and Drp1 interaction mutants, *cnc1Δ* cells harboring the expression constructs described in Fig 5A were treated with RH123 before and following CHP exposure. These studies revealed that MOMP was observed in the Cdk8-interaction mutants consistent with the normal cell death response. However, MOMP was not observed with the Drp1-interaction mutations. These results indicate that Cnc1-Drp1 interaction is required for cell death at either the execution of MOMP or at a step prior to this process.

### The human cyclin C induces mitochondrial fission and supports RCD in yeast

Previous studies (38), including our work (39), found that CCNC could not complement the transcription defect in *cnc1Δ* mutant yeast. However, recombinant Cnc1 induced mitochondrial fission when added to permeabilized mouse embryonic fibroblasts (13). To test whether CCNC can induce mitochondrial fission and/or RCD in yeast, a single-copy plasmid harboring an EGFP-*CCNC* reporter gene expressed from the *ADH1* promoter was transformed into a *cnc1Δ* strain. Using the nuclear pore complex component Nup49 (40) fused to mCherry (a gift from K. Madura), fluorescence microscopy revealed that EGFP-CCNC localizes to the nucleus in unstressed cells (Fig 6A, see Fig S6A for field view). Next, we repeated these experiments with the *cnc1Δ* strain expressing EGFP-CCNC and a mitochondrial targeted DsRed reporter protein. In CHP treated cells, EGFP-CCNC re-localized to the cytoplasm where it associated with fragmented mitochondria (arrows, Fig 6B, see Fig S6B for field view). These results indicate that the CCNC efficiently translocates from the nucleus to the mitochondrial and stimulates fragmentation in yeast.

**Figure 6.**
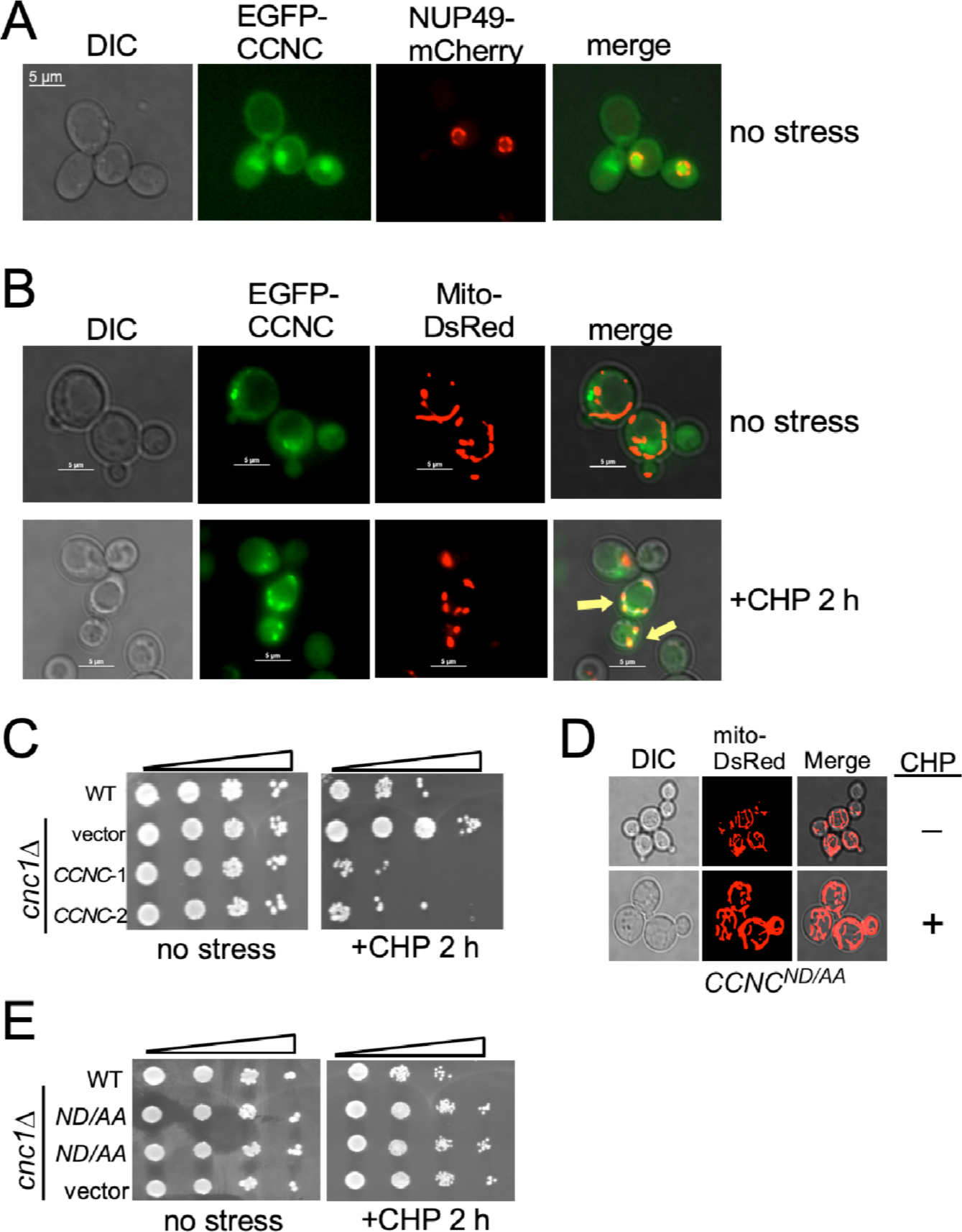
Human CCNC induces fission and supports RCD in yeast. Fluorescence microscopy imaging mitochondrial morphology (red) and EGFP-CCNC (green) localization before (A) and following (B) CHP treatment (50 µM). Arrows indicate colocalization of EGFP-CCNC and mitochondria in stressed images. (C) Viability studies were conducted as described in Fig 5. Two independent transformants expressing EGFP-CCNC were assayed. Predicted DRP1 interaction sites (N181, D182) are required for CHP-induced mitochondrial fission (D) and RCD (E). Two independent transformants were assayed for RCD following CHP treatment.

Next, the ability of EGFP-CCNC to mediate RCD in yeast was tested as described earlier. Two independent transformants expressing EGFP-CCNC were grown to mid-log phase then treated with CHP for 2 h. These samples, and those without treatment, were serially diluted (1:5) then plated on complete minimal medium. As expected, the wild type and *cnc1Δ* strains exhibited sensitivity and resistance to CHP, respectively (Fig 6C). Interestingly, EGFP-CCNC expression induced RCD with efficiencies equal to the wild-type yeast. To determine if the presence of the EGFP moiety affected these results, the experiments were repeated with a *CCNC* cDNA expression construct again expressed from the *ADH1* promoter. These results revealed a similar sensitivity to CHP as observed with the EGFP-tagged CCNC (Fig S6B). Finally, we tested whether overexpression of CCNC would affect overall cell viability as well as CHP sensitivity. Expressing *ADH1pro*-*CCNC* from a high-copy 2µ plasmid did not affect overall cell viability but still supported CHP sensitivity. Finally, we determined whether CCNC was inducing cell death via Dnm1 interaction by introducing a double substitution mutation (ND181,182AA) in a predicted DRP1 interaction region. This derivative was expressed at the same level as wild type (Fig 6D) indicating that the mutations do not alter protein stability. Finally, this mutant was resistant to CHP treatment suggesting that, similar to Cnc1, CCNC induces cell death in yeast through Dnm1 interaction. These results indicate that CCNC can perform both the mitochondrial fragmentation and cell death functions in yeast. In addition, these results indicate an important role for CB2 in mediating cell death that is conserved from yeast to human.

## Discussion

Cyclin C is a highly conserved component of the Mediator Kinase Module (MKM) that directly controls transcription through stimulation of its cognate kinase CDK8. In response to oxidative stress, cyclin C also translocates to the mitochondria where it binds and stimulates the fission factor DRP1 or Dnm1. We previously demonstrated that cyclin C is required for oxidative stress-induced regulated cell death (RCD) in both yeast and mammalian cells. Therefore, the role of cyclin C in RCD could depend on its transcriptional role, mitochondrial activity, or both. To address this question, we utilized protein-protein binding algorithms that identified CCNC-CDK8 and CCNC-DRP1 interaction sites in cyclin box 1 (CB1) or cyclin box 2 (CB2), respectively. Our studies revealed that association with the fission factor Dnm1, but not Cdk8, was required for RCD in yeast. However, although Dnm1 was required for RCD, its fission function was not. In support of this conclusion, the human cyclin C (CCNC) that does not regulate transcription in yeast but can efficiently induce mitochondrial fission and RCD. Moreover, CCNC appears to utilize the same pathway as this activity is dependent on its interaction with Dnm1. Taken together, this study finds that the mitochondrial role of cyclin C is critical for RCD in yeast and demonstrates the remarkable conservation of CB2 function with the human and yeast proteins.

Our findings highlight three important considerations when using docking modeling tools such as ClusPro. First, all protein-protein docking simulators have some inherent error and uncertainty associated with their use. For example, rigid body modeling has limitations as modeled lack side-chain flexibility, molecular relaxation, or conformational changes (41, 42). Secondly, while modeled interacting residues can show a moderate degree of overlap with true experimentally derived residues, as we observed for our modeling of CCNC-CDK8, this does not necessarily mean the entire structural similarity overlaps. Lastly, one must be careful when using RMSD calculations to grade structural alignments. A broad range of RMSD values were observed when aligning modeled with experimentally derived complexes, despite quality identification of interacting residues. Using the default settings of PyMol’s built in alignment tool, with RMSD distance cutoffs set to 2 Å. This resulted in nearly 90% of models having a RMSD value less than 3.0 Å. Extending the cutoff to only 3 Å resulted in only 20% of models having a RMSD value less than 3.0 Å. This highlights important considerations that are needed when using RMSD analyses tools to overlay protein structures. However, our genetic results indicated that >90% of the predicted interactions were important for protein function. Therefore, this approach can provide valuable data for identifying function protein-protein interactions.

We identified Cnc1 amino acids whose mutation was predicted to disrupt the interaction with either CDK8 or DRP1. Our hypothesis predicted that these mutations should reduce protein interaction causing a loss in either transcriptional repression or mitochondrial fission function, respectively. This model proved correct in that fission mutations reduced, although did not eliminate, Dnm1-Cnc1 association. However, no difference in co-immunoprecipitation efficiency was observed between Cnc1 transcription mutants and Cdk8 compared to the control. These differences may be due to the context in which these proteins interact. Cnc1 and Cdk8 associate with the large scaffolding proteins Med12 and Med13 forming the highly conserved MKM (43). For example, Cnc1 interacts with Med13 through an amino terminal alpha helix (26). Using a genetic approach, we also identified this domain which we termed the holoenzyme association domain (HAD, see Fig 1C) (44) that associates with Med13 in both yeast (45) and mice (46). In the other direction, haplo-insufficiency of *MED13L* in the developmental intellectual disability *MED13L* syndrome results in aberrant CCNC nuclear release and constitutive mitochondrial fragmentation and dysfunction (47). These results revealed that the scaffold attachments of the MKM are highly conserved and play a critical role in maintaining the integrity of this complex.

How does cyclin C induce cell death in yeast? In this study, we found a 100% correlation between RCD execution and MOMP. In mammals, MOMP generally represents the point of no return as release of cytochrome *c* triggers caspase activation necessary for cellular dismemberment. In addition, MOMP releases additional pro-apoptotic proteins such as the nuclease EndoG (48) and the apoptotic inducing factor AIF (49). However, a role for MOMP and cytochrome *c* release in RCD execution in invertebrates and fungi is unclear. For example, even though caspases exist in *Drosophila* and *C. elegans*, cytochrome *c* release is not necessary for their activation (50–52). Similarly, cytochrome c release in yeast is not required for activation of the yeast metacaspase Yca1 (53). Interestingly, release of Nuc1, a pro-apoptotic homolog of EndoG (54) and Aif1 (55) is also observed. However, deleting these genes only partially protected cells from ROS stress. This contrasts with *cnc1Δ* or *dnm1Δ* mutants which provide dramatic protection (see Fig 5A for example). These results suggest that, similar to mammalian cells, MOMP is the key regulatory step in RCD execution. Therefore, these results suggest a model that yeast RCD occurs through release of multiple cell death factors stemming from loss of mitochondrial integrity. Alternatively, there may be another, more universal mode of cell death, that is yet to be described in yeast.

The cyclin family is defined by the presence of a cyclin box (CB1) in the amino terminal portion of the protein that binds the Cdk. CB1 is highly conserved among all cyclin families as it is constrained to mediate CDK binding. In addition, a second carboxyl located cyclin box (CB2) is observed in both canonical and transcription cyclins but not for a large group of “atypical” cyclin proteins (56). Our results revealed that both the yeast and human cyclin C CB2 are critical for mitochondrial fission and RCD in yeast. Interestingly, this region was identified in a structural study as containing an acidic cleft potentially binding proteins other than Cdk8 (18). Although CB2 is relatively well conserved within an individual cyclin subfamily, it becomes very divergent when compared with other groups. For example, helix H2’ and H3’ is highly conserved with C-type cyclins from *S. pombe*, *S. cerevisiae* and *H. sapiens* (18). However, they are divergent from cyclin H or cyclin A family members. These findings suggest that the duplication of the cyclin box domain allowed individual proteins to acquire new activities through divergent processes. These findings suggest that other cyclin subfamilies that maintain conserved CB2 domains may also have an additional evolutionarily conserved function.

## Methods

### Protein structural analysis and modeling

Structure files (.pdb) of experimentally solved structures were accessed on the RCSB PDB (57). For proteins without known structures, models were generated using either AlphaFold (24) or SWISS-MODEL (58). Protein docking simulations were run on the ClusPro 2.0 server (21, 22) with default configuration settings. Protein-protein docking simulations under *Balanced*, *Electrostatic*, and *Hydrophobic* ClusPro classifications were exported and edited using PDB Editor (59) or PDB-tools (22) to modify chain IDs for downstream analysis. Experimentally solved or simulation-generated protein-protein structures were uploaded to the PDBePISA (Protein, Interfaces, Structures, and Assemblies) online suite (60), allowing for identification of interaction interfaces and interacting residues between modeled chains. PISA results from all generated protein-protein models were compiled, adjusted by removing impossible interactions or those deemed an artifact of analysis, and ranked by the frequency of both residue pairs and individually highlighted CCNC residues.

Visualization of protein structural files, simulated models, and protein complexes was accomplished in PyMol (19). To analyze structural differences in three-dimensional space between two or more protein or complex structures, PyMol’s built in alignment plugin was used with settings adjusted to ten cycles with a 2.0 cutoff. The output of this analysis is the RMSD value, or a numerical measurement representing the average deviation between corresponding atoms of two structures (61). Protein sequence comparison was performed using both local (62) and global (63) protein sequence alignment services from the NCBI BLAST online suite. Additional tools such as PrDos (64), IUPred2 (65), and flDPnn (66) were used for analysis and mapping of intrinsically disordered domains.

### Yeast strains and plasmids

Unless otherwise noted, experiments were performed in the *S. cerevisiae* W303 background RSY10 (*MAT***a** *ade2 ade6 can1-100 his3-11,15 leu2-3,112 trp1-1 ura3-1*) and the following derivatives RSY391 (*cnc1*::*TRP1*) and RSY1696 (*cnc1*::KanMX) (67). In accordance with the Mediator nomenclature unification efforts (43), yeast cyclin C (*SSN8/UME3/SRB11/CNC1*) will use the *CNC1* gene designation. Plasmids used in this study are listed in Table S3. The Mt-DsRed expression plasmid was previously described (68). Site directed mutagenesis was accomplished using PCR with Phusion^TM^ High-Fidelity DNA Polymerase (ThermoFisher #F530S) and SDM primers (see Table S2) generated with NEBaseChanger (New England Biolags #E0554S). All mutations were verified by DNA sequence analysis.

### Yeast viability assays

Yeast strains were grown in selective synthetic minimal dextrose (SD) medium (0.17% yeast nitrogen base without amino acids and ammonium sulfate, 0.5% ammonium sulfate and 2% glucose supplemented with amino acids/uracil to allow for plasmid selection (69). For all experiments, the cells were grown to mid-log phase (∼6 × 10E6 cells/mL) before directly adding 50 µM cumene hydroperoxide (CHP) for 2 h. Untreated cells were grown for an additional 2 h to serve as controls for treated cells. Both treated and untreated cells were then spotted on complete SD plates using 5-fold serial dilutions. Images of the plates were captured using an iBright FL1500 imaging system following 2 d incubation at 30°C.

### RT-qPCR Analysis

Total RNA was prepared using the Monarch^®^ Total RNA Miniprep Kit and glass bead lysis. On-column DNase treatment was performed to eliminate contaminating DNA during RNA extraction. Total RNA (1 μg) was converted to cDNA using the Maxima cDNA Synthesis kit (ThermoFisher #K1671). The cDNA from each sample was diluted (1/100) and subjected to qPCR amplification using PowerSYBR^®^ Green PCR Master Mix (ThermoFisher #4367659) and a StepOne^TM^ Real Time PCR System (ThermoFisher). These assays were conducted with three independent samples assayed in triplicate. *ACT1* was used as the internal standard for comparative (ΔΔC_T_) quantification. Statistical significance was determined via Student’s T-test analysis. Primer sequences for RT-qPCR analysis can be found in Table S3.

### Fluorescence Microscopy/Mitochondrial Dynamics Quantitation

For all microscopy studies, cells were grown to mid-log phase prior to CHP treatment. Deconvolved images were obtained using a Nikon microscope (Model E800) with a 100x objective with 1.2x camera magnification (Plan Fluor Oil, NA 1.3) and a CCD camera (Hamamatsu Model C4742). Data were collected using NIS software and processed using Image Pro software. All images of individual cells were optically sectioned (0.3-μM spacing), deconvolved and slices were collapsed to visualize the entire fluorescent signal within the cell. The mitochondria were visualized by expressing a mitochondrial targeted DS-red construct or staining with MitoTracker Red CMXRos (ThermoFisher #M7512). Yeast and human cyclin C were visualized using YFP or EGFP fluorescence. Mitochondrial morphology was scored reticular if 1-3 long filaments were observed. Fragmented phenotype ≥5 short tubules were noted.

### Western blot analysis/Quantitative co-immunoprecipitation assays

Soluble extracts were prepared using glass bead lysis of mid log phase cells. 50 µg of lysate were separated by PAGE, blotted to Immobilon-P nylon membrane (MilliporeSigma #IPVH10100) the probed with αCCNC antibodies (Thermo Fisher #PA5-16227). For co-immunoprecipitation studies, cells were grown to mid-log phase (5-7×10E6/ml), treated with 50 µM CHP for 4 h, then harvested by centrifugation. Cell pellets were either frozen or directly lysed using glass bead protocol (70). One mg of soluble protein was incubated with 10 µL of ChromeTek GFP-trap^®^ beads (Proteintech #Gta-20.) for one h at 4°C. The beads were extensively washed, mixed with sample buffer +DTT and separated by electrophoresis on a 10% acrylamide gel. The gel was subjected to Western blot analysis and specific proteins visualized using mouse αmyc (EMD Millipore #05-724) or αHA (Abcam Cat# ab9110, RRID:AB_307019) primary antibodies. Pgk1 levels were monitored using rabbit αPgk1 antibodies (ThermoFisher, #459250). These signals were quantified using an alkaline-phosphatase-conjugated goat anti-mouse (Abcam Cat# ab97027, RRID:AB_10679837) or anti-rabbit (Abcam Cat# ab97061, RRID:AB_10680575) secondary antibodies using the CDP-Star chemiluminescence kit (Invitrogen, #T2307). Two quantify the signals, the blots were exposed on the 1500 iBright Chemiluminescence Imager (Invitrogen). These values were then normalized to the Pgk1 loading control signals. Cnc1/Dnm1 or Cnc1/Cdk8 ratios were calculated following normalization with WT values set to 1.0. P-values shown are relative to wild-type 4 h timepoints.

### Statistical analysis

All representative results included at least two independent biological experiments. P values were generated from Prism-GraphPad using unpaired Student’s *t*-tests; NS P ≥ 0.05; *P ≤ 0.05, **P ≤ 0.005; ***P ≤ 0.001; ****P ≤ 0.0001. All error bars indicate mean ± SD.

## Acknowledgments

We thank K.F. Cooper and V. Ganesan for plasmids and helpful comments. We thank K.F. Cooper and B. Wiser for critical reading of this manuscript. We thank J. Nunnari for the Mt-DsRed expression plasmid.

## AUTHOR CONTRIBUTIONS

S.J.D, Investigation, Formal analysis, writing draft; J.R.B., Investigation; Formal analysis, Writing draft; D.C.S., Investigation, Formal analysis; S.E., Investigation; D.G.J.S., Investigation; R.S. Draft Review, Editing, Supervision, Funding acquisition, Resources.

## Supporting information

Table S1. Plasmids used in this study.

Table S2. Primers used in this study for site directed mutagenesis.

Table S3. Primers used in this study for RT-qPCR analyses.

Figure S1. Analysis of yeast and human protein homology.

Figure S2. Structural analysis of DRP1.

Figure S3. Mapping cyclin C residues predicted to interact with CDK8 or DRP1.

Figure S4. Quantitation of Cnc1-Dnm1 co-immunoprecipitation.

Figure S5. Cnc1-dependent MOMP induction in response CHP treatment.

Figure S6. Human CCNC induces mitochondrial fragmentation and promotes RCD in yeast.

